# Melatonin Partially Attenuates Oxycodone-Induced Placental Stress Signaling and Fetal Brain Apoptosis in a Sex-Specific Manner

**DOI:** 10.64898/2026.04.29.721662

**Authors:** IO Adediji, K Kamra, HM Kowash, P Nouri Mousa, CO Aloba, VL Schaal, JS Davis, ES Peeples, G Pendyala, LK Harris

## Abstract

**Background:** Maternal oxycodone (oxy) exposure can disrupt placental function and fetal neurodevelopment, but the molecular mechanisms remain unclear. We investigated whether prenatal oxy exposure activates inflammation and stress response pathways in the placenta and fetal brain, and if maternal melatonin supplementation attenuates these effects.

**Methods:** Female Sprague-Dawley rats received either saline or oxy via oral gavage for 15 days before mating (10-15mg/kg/day dose escalation) and throughout pregnancy (15mg/kg/day). From gestational day (GD) 12.5, half of the dams received melatonin (10mg/kg/day). On GD 19.5, placental and fetal brain tissues were collected. Changes in expression of markers of oxidative stress, antioxidant defense signaling, inflammation, ER stress, and apoptosis were assessed by western blotting. Data were analyzed by two-way ANOVA with Tukey’s post hoc test.

**Results:** Neither oxy exposure nor melatonin treatment increased markers of oxidative stress or antioxidant defenses in the placenta and fetal brain. Oxy exposure increased placental IL-1β expression but did not alter expression of the other inflammatory markers examined. Oxy increased phosphorylation of eIF2α and increased the phospho-eIF2α:eIF2α ratio in the placentas of male fetuses, and fetal brains of both sexes. CHOP expression was increased in the placentas and brains of female, but not male fetuses after oxy exposure. Oxy exposure increased levels of cleaved caspase-3 and cleaved caspase-9 in the fetal brain, but not the placenta; melatonin treatment attenuated the oxy-induced increase in cleaved caspase-9, but not cleaved caspase-3.

**Conclusion:** Prenatal oxy exposure induced a modest inflammatory response in the placenta and activated the integrated stress response and intrinsic apoptotic signaling in the fetal brain. Maternal melatonin supplementation partially mitigated the oxy-induced upregulation of caspase-9 but did not prevent stress signaling in either tissue. These findings demonstrate the presence of sex-specific placental and fetal brain responses to prenatal oxy exposure but suggest that melatonin may not provide complete protection against oxy-induced neurodevelopmental impairment.

## INTRODUCTION

Prenatal opioid exposure remains a significant public health challenge, with consequences extending beyond neonatal withdrawal at delivery, to include placental dysfunction and neurodevelopmental impairment^1^. Among prescription opioids, oxycodone (oxy) use in pregnancy is of particular concern, due to well-documented evidence of fetal exposure^2^. Oxy also exhibits a higher rate of placental transfer compared to other commonly used opioids^2^. The placenta is a dynamic mediator of fetal development, and emerging evidence suggests that opioid-induced alterations in placental development and function may directly or indirectly influence fetal brain development^3, 4^.

Prenatal oxy exposure has been shown to disrupt the structural integrity of the placental labyrinth and alter trophoblast differentiation in mice, processes critical for adequate placental transport capacity^4,5^. In addition to these structural alterations, studies in rodents have shown that oxy exposure impairs placental vascularization, perfusion, and nutrient transport, providing additional routes through which opioid exposure may affect fetal growth and developmental trajectory^6-9^. High-throughput transcriptomic profiling in oxy-exposed rodents has shown that oxy directly alters stress-responsive signaling pathways in the developing brain, including glutamatergic, synaptic transmission, and reward-related gene networks in the prefrontal cortex and nucleus accumbens, suggesting these neurological disruptions may represent early mechanistic links between maternal oxy exposure and downstream neurological injury^10-14^.

Chronic opioid exposure has been linked to oxidative stress in differentiated human SH-SY5Y neuroblastoma cells^15, 16^, in the liver and hippocampus of rodents^17, 18^. It has also been associated with dysregulated inflammatory cytokine profiles in cancer patients on long-term opioid therapy^19^ and the activation of integrated stress response (ISR) in the brain of female rats chronically treated with oxy^14^. Oxidative stress involves the accumulation of excess reactive oxygen species (ROS), which drive lipid peroxidation and generate reactive aldehydes such as 4-hydroxynonenal (4-HNE) and malondialdehyde (MDA), which form covalent adducts with proteins and nucleic acids^20^. Accumulating evidence shows that prenatal opioid exposure in rodents increases lipid peroxidation in neonatal tissues^17, 21, 22^, including elevated protein carbonyls^22^.

While physiological levels of ROS act as signaling molecules for trophoblast differentiation and angiogenesis^23, 24^, excessive oxidative stress is a major contributor to placental dysfunction^25^. To counteract the deleterious effects of ROS on placenta function and fetal growth, the placenta upregulates its own endogenous antioxidant mechanisms, particularly the Kelch-like ECH-associated protein 1 (KEAP1)-Nuclear factor erythroid 2-related factor 2 (NRF2) signaling pathway, which regulates the cellular antioxidant response^23, 24, 26^. Mechanistically, KEAP1 usually binds NRF2 in the cytoplasm and targets it for proteasomal degradation, but under prolonged oxidative stress, KEAP1 cysteines are modified, NRF2 is stabilized and translocates to the nucleus, where it binds to antioxidant response elements (AREs) to drive transcription of antioxidant genes like heme-oxygenase-1 (HMOX1)^24, 27^. Opioids like morphine induce oxidative stress in tissues and dysregulate the KEAP1/NRF2 pathway in a context-dependent manner. In primary human brain microvascular endothelial cells, morphine upregulated several NRF2 target enzymes, including HMOX1 and thioredoxin reductase 1 (TXNRD1), while simultaneously decreasing catalase and the NRF2 upstream regulator protein/nucleic acid deglycase (DJ-1)^28^. In contrast, morphine suppressed the NRF2 signaling in osteoblasts, reducing the expression of HMOX1 and TXNRD1^29^, demonstrating that the downstream antioxidant response to opioid-induced oxidative stress is highly cell type-specific.

Beyond direct oxidative injury, ROS amplify inflammatory signaling by activating pattern recognition receptors like Toll-like receptor 4 (TLR4) and upregulating pro-inflammatory cytokines including interleukin-6 (IL-6), IL-1β, and tumor necrosis factor-α (TNF-α). These inflammatory mediators in turn trigger endoplasmic reticulum (ER) stress and the ISR, creating a positive feedback loop, which is particularly common in tissues with high secretory demand, like the placenta^30, 31^. The ER is the principal site of protein folding and is highly sensitive to the metabolic stressors imposed by prenatal drug exposure^30, 31^. When folding demand exceeds capacity, the unfolded protein response (UPR) is activated and triggers the ISR, a conserved homeostatic program^32^. This in turn causes one or more of the four upstream kinases central to ISR activation, including protein kinase R-like endoplasmic reticulum kinase (PERK; activated by ER stress), heme-regulated inhibitor (HRI; activated by heme deprivation and oxidative stress), general control non-derepressible 2 (GCN2; activated by amino acid deprivation), and protein kinase R (PKR; activated by double-stranded RNA and inflammatory signals), to phosphorylate eukaryotic initiation factor 2α (eIF2α) at Ser51 and suppress cap-dependent translation, while paradoxically allowing the translation of activating transcription factor 4 (ATF4)^32, 33^. The cytokine-driven ROS production and ER calcium dysregulation downstream of NF-κB can activate PERK and HRI, providing a mechanistic bridge from inflammation to ISR activation^30, 34^. ATF4 in turn drives transcription of C/EBP homologous protein (CHOP), which switches the response from adaptation toward apoptosis by repressing the anti-apoptotic protein Bcl-2 while upregulating pro-apoptotic factors including Bcl-2-interacting mediator of cell death (Bim), and Death receptor 5 (DR5)^30, 34^. These changes converge on the mitochondrial outer membrane, increasing its permeability and triggering the cleavage and activation of caspase-9 and caspase-3^30, 34^.

Recent studies have shown sustained PERK-eIF2α-ATF4-CHOP signaling in preclinical models of gestational stress, including maternal nicotine exposure, where it is associated with placental insufficiency, altered trophoblast function, and fetal growth restriction^32, 35^. Similarly, prenatal opioid exposure has been linked to placental dysfunction and adverse fetal brain outcomes in preclinical models^4, 10-13^,however whether these coordinated stress responses are directly activated in the placenta and fetal brain in the context of prenatal oxy exposure remains less clear.

We have previously established a rat model of prenatal oxy exposure that exhibited fetal growth restriction and neurodevelopmental abnormalities in exposed offspring^10-13^, supporting the premise that placental dysfunction and fetal brain injury may be mechanistically linked. Despite the documented harm of prenatal opioid exposure, there is currently no medication available to mitigate placental and fetal brain injury in this setting. Melatonin has emerged as a potential candidate, given that, in addition to its role as a regulator of circadian rhythms, it is a potent antioxidant and anti-inflammatory agent that readily crosses the placenta^36^. In rodent models of fetal growth restriction, melatonin has been shown to attenuate placental endoplasmic reticulum stress and reduce trophoblast apoptosis^37^. Melatonin has also been shown to reduce placental oxidative stress, inflammation, mitochondrial dysfunction, and fetal brain injury in preclinical models^38-40^. Therefore, in this study, we investigated whether prenatal oxy exposure in pregnant rats caused oxidative stress, activated antioxidant response signaling, induced inflammation or triggered ER stress in the placenta and fetal brain. We further examined whether exogenous melatonin supplementation mitigated oxy-induced alterations and if any of the observed responses was sex dependent.

## MATERIALS AND METHODS

### Ethical Approval

Animals were housed in a temperature-controlled environment (22 °C – 25 °C) with a 12-hour light–dark cycle and *ad libitum* access to food and water, in accordance with standards set by the National Institutes of Health Guidelines for the Care and Use of Laboratory Animals^41^. All experimental protocols were approved by the Institutional Animal Care and Use Committee (IACUC) of the University of Nebraska Medical Center (protocol ID no. 17-080-09 FC).

### Animals

Male (n = 10) and female Sprague Dawley rats (n = 30) were purchased from Charles River Laboratories Inc. (Wilmington, MA, USA). Animals were housed on-site and allowed a 1-week acclimation prior to experimentation. Delivery of drugs/chemicals (saline, oxy and /or melatonin) was administered within our animal housing center. All other experiments were conducted in our basic science lab.

### Experimental Design

Oxycodone HCl (Sigma Aldrich, St. Louis, MO, USA) exposure was adapted from our previously published studies^10-13^. The melatonin dosing regimen was based on previously published work showing significant improvements in placental function and neonatal outcomes^42-44^. Briefly, between 8 - 9 am each morning, nulliparous female rats (64 - 70 days of age) received either saline (sal) or oxy by oral gavage for 5 days, starting with a dose of 10 mg/kg/day, which was then increased by 0.5 mg/kg/day over the next 10 days to reach a final dose of 15 mg/kg/day; on day 15, the female rats were mated with proven male breeders. Pregnancy was confirmed by the presence of a vaginal plug the following morning, and the day of plugging was designated as gestational day (GD) 0.5. This treatment regimen continued throughout gestation; however, from GD12.5 to GD19.5, half of the dams in each group additionally received oral melatonin (Sigma Chemical Co., St. Louis, MO, USA) at a dose of 10 mg/kg/day, resulting in four experimental groups: saline (Sal), saline and melatonin (Mel), Oxy, and Oxy and melatonin (OxMe).

### Tissue Harvest

On GD19.5, dams were euthanized by isoflurane overdose (5%; Abbott, UK) one hour after their final treatment. Maternal blood was collected by cardiac puncture and centrifuged at 1000 x g at 4 ◦C for 10 minutes; plasma was aliquoted into sterile tubes and snap frozen. The uterine horn was removed and placed on ice; fetuses and placentas were removed and weighed. Maternal and fetal organs were then dissected, weighed and either snap frozen, stored in RNA later or fixed for histological examination. Fetal tail tips were collected for sex determination, performed by PCR amplification of the male-specific Sry gene, as previously described^45, 46^.

### Western Blot Analysis

Western blotting was performed to determine the relative protein expression of oxidative stress, ER stress, inflammatory, and apoptotic markers in placental and fetal brain tissues. Frozen placenta and fetal brain tissues were homogenized in ice-cold lysis buffer prepared from RIPA buffer supplemented with Halt Protease and Phosphatase Inhibitor Cocktail (Thermo Scientific, #1861281) and centrifuged, and supernatants were collected for the determination of protein concentration. Protein concentration was determined using the Pierce BCA Protein Assay Kit (Thermo Scientific, #23225) and compared against a standard curve prepared with bovine serum albumin. Equal amounts of protein were resolved on NuPAGE Bis-Tris Mini Protein Gels (4 - 12 %, NP0323BOX) and transferred onto polyvinylidene fluoride (PVDF) membranes (Thermo Scientific, #88520).

Membranes were blocked with 5% non-fat dry milk in 1x TBST containing 0.1%(v/v) TWEEN 20 (Sigma-Aldrich, Cas #9005-64-5), for 1 hour at room temperature and incubated overnight at 4°C with primary antibodies against NRF2 (1:500, Cell Signaling Technology, CST#336495), phospho-NRF2 (1:500, Sigma-Aldrich, SAB#5701092), KEAP1 (1:1000, Cell Signaling Technology, CST#8047), HMOX1 (1:1000, Cell Signaling Technology, CST#43966), phospho-eIF2α (Ser51; 1:500, Cell Signaling Technology, CST#3398), eIF2α (1:500, Cell Signaling Technology, CST#5324), BiP (GRP78) (1:500, Cell Signaling Technology, CST#3183), CHOP (1:500, Cell Signaling Technology, CST#2895), ATF4 (1:500, Cell Signaling Technology, CST#11815), TLR4 (1:500, Abcam, #ab22048), IL-1β (1:500, Abcam, #ab254360), TNF-α (1:500, Abcam, #ab215188), cleaved caspase-3 (1:1000, Cell Signaling Technology, CST#9664), cleaved caspase-8 (1:500, Cell Signaling Technology, CST#9429), cleaved caspase-9 (1:500, Cell Signaling Technology, CST#9507), MDA (1:1000, Abcam, #ab243066), 4-HNE (1:500, Abcam, #ab48506), and GAPDH (1:1000, Cell Signaling Technology, CST#2118) as a loading control. Primary antibodies were dissolved in 5% BSA (Fisher bioreagents, BP1600-100) in 1x TBST.

Following washing with 1x TBST three times for 5 minutes each, membranes were incubated with horseradish peroxidase-conjugated goat anti-rabbit IgG secondary antibody (1:10,000, Jackson ImmunoResearch, #111-035-045) or goat anti-mouse IgG secondary antibody (1:10,000, Jackson ImmunoResearch, #115-035-205), as appropriate, for 1 hour at room temperature, followed by 3 washes for 5 minutes each with 1x TBST. Protein bands were visualized using Sensitivity Chemiluminescent Substrate (Thermo Scientific, #A38556) and imaged using iBright imaging system (Invitrogen Thermo Fisher Scientific CL 1500). Densitometric analysis was performed using ImageJ (v1.54p, NIH) and protein expression was normalized to GAPDH and expressed as a percentage of control gene expression.

### Statistical Analysis

Data analysis in text and figures is presented as mean ± SEM. Statistical evaluation was analyzed using GraphPad Prism v10.6 (GraphPad Software, San Diego, CA, USA). Comparisons between groups utilized two-way ANOVA followed by Tukey’s post hoc test with P < 0.05 considered statistically significant.

## RESULTS

### Prenatal oxy exposure does not alter expression of oxidative stress and antioxidant response markers in the placenta or fetal brain

To assess whether prenatal oxy exposure would induce oxidative stress and upregulate antioxidant response genes in the placenta or fetal brain, we measured the expression of 4-HNE and MDA-modified proteins, as well as NRF2, KEAP-1, and HMOX-1, at GD19.5 of pregnancy. Neither oxy exposure, nor melatonin treatment altered the expression of any of these markers in either the placenta or fetal brain, and no sex-dependent effects were observed (Figures 1 and 2). These findings indicate that oxy exposure, with or without melatonin treatment, does not induce significant levels of oxidative stress or upregulate antioxidant defense pathways in either tissue at this gestational time point in our rat model.

**Figure 1.**
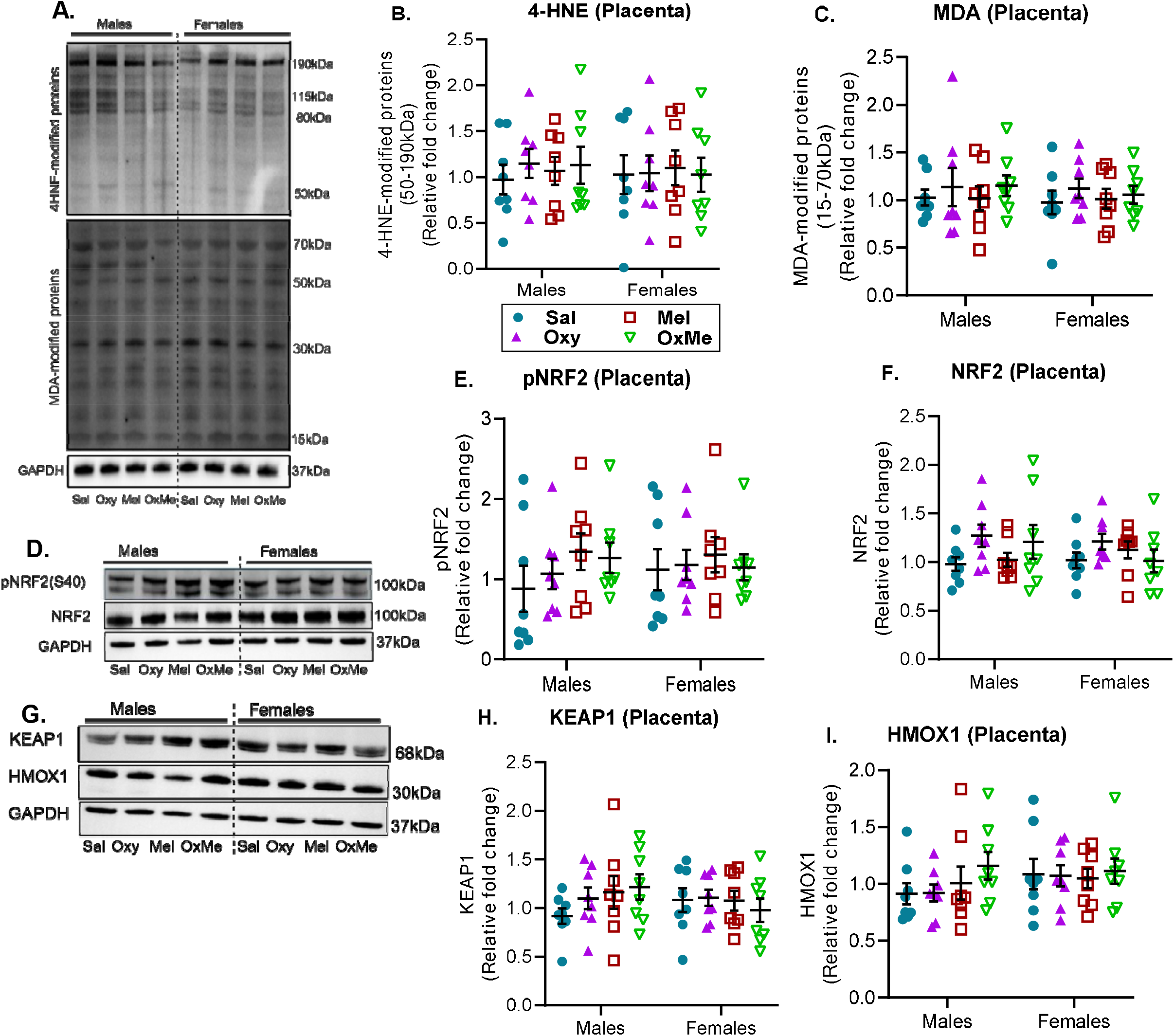
Effects of oxy and melatonin on oxidative stress and antioxidant response markers in GD19.5 placenta. (A-C) Representative western blot images and densitometric quantification of 4-HNE and MDA protein adducts. (D-F) Representative western blot images and densitometric quantification of phosphoNRF2 and NRF2. (G-I) Representative western blot images and densitometric quantification of KEAP1 and HMOX1. All samples were normalized to GAPDH. Data were analyzed by a 2-way ANOVA followed by Tukey’s post hoc test and presented as mean ± SEM. n=8/sex/group. *P*<0.05 was considered statistically significant.

**Figure 2.**
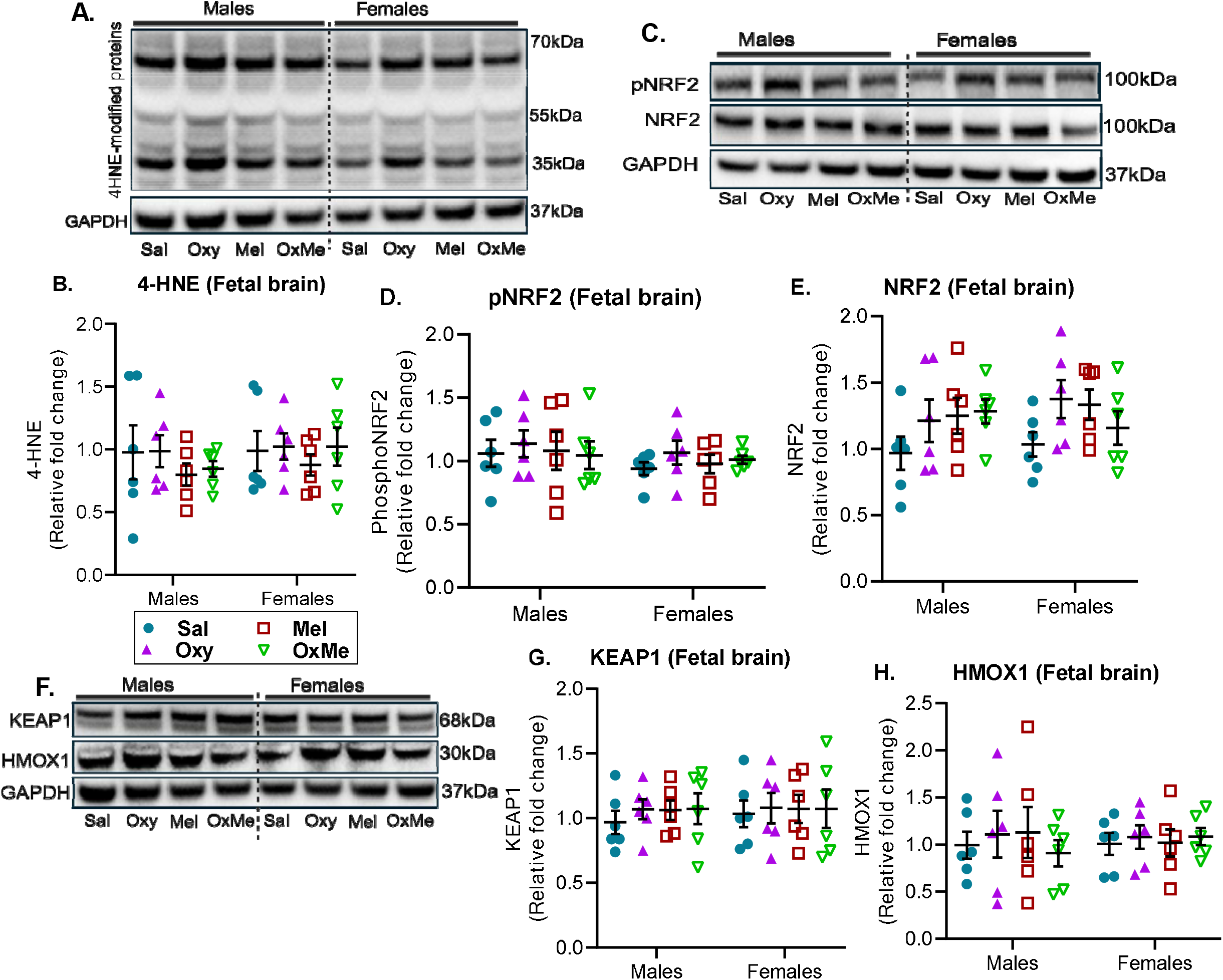
Effects of oxy and melatonin on oxidative stress and antioxidant response markers in GD19.5 fetal brain. (A-B) Representative western blot images and densitometric quantification of 4-HNE protein adducts. (C-E) Representative western blot images and densitometric quantification of phosphoNRF2 and NRF2. (F-H) Representative western blot images and densitometric quantification of KEAP1 and HMOX1. All samples were normalized to GAPDH. Data were analyzed by a 2-way ANOVA followed by Tukey’s post hoc test and presented as mean ± SEM. n=6/sex/group. *P*<0.05 was considered statistically significant.

### Prenatal oxy exposure increases placental expression of IL-1β, **but not TLR4 or TNF-**α

To characterize the inflammatory profile of the placenta following prenatal oxy exposure, we measured protein levels of IL-1β, TNF-α, and TLR4. Placental expression of TLR4 and TNF-α was unchanged across all the groups (Figure 3A-E); however, expression of IL-1β was significantly elevated by oxy in placentas from both male (p=0.011; Figure 3B) and female fetuses (p=0.013; Fig. 3B). Melatonin co-treatment caused a 40% decrease in the mean level of IL-1β expression, but did not reach the levels seen in the controls (Figure 3B).

**Figure 3.**
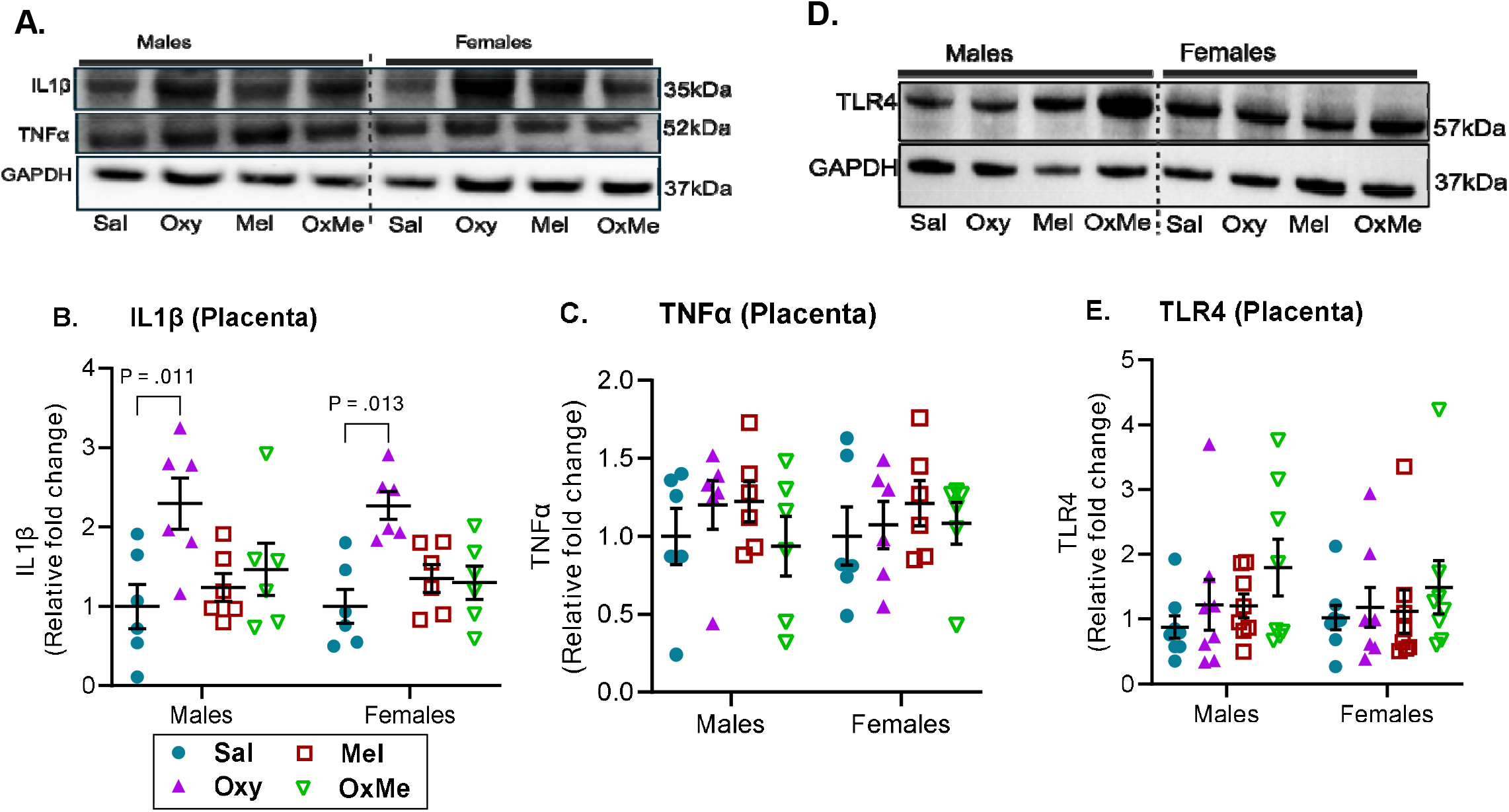
Effects of oxy and melatonin on selected inflammatory markers in GD19.5 placenta. (A-C) Representative western blot images and densitometric quantification of IL1β and TNFa. (D-E) Representative western blot images and densitometric quantification of TLR4. All samples were normalized to GAPDH. Data were analyzed by a 2-way ANOVA followed by Tukey’s post hoc test and presented as mean ± SEM. n=6/sex/group. *P*<0.05 was considered statistically significant.

### Prenatal oxy exposure activates the integrated stress response, but not the canonical ER stress pathway in the placenta and fetal brain

To investigate whether prenatal oxy exposure can activate the ER stress pathway, we assessed expression levels of BiP, total eIF2α, phosphorylated eIF2α (p-eIF2), ATF4 and CHOP in both the placenta and fetal brain. Expression of ATF4 and BiP was unaffected by oxy exposure or melatonin treatment in the placenta (Figure 4A-C) and the fetal brain (Figure 5A-C).. Also, levels of total eIF2α were unaffected by either oxy exposure or melatonin treatment in both placenta and fetal brain (Figures 4D and 5D). Together, these results indicate that the canonical ER stress was not activated under these experimental conditions. In contrast, p-eIF2α levels and the p-eIF2α:eIF2α ratio were significantly elevated in oxy-exposed placentas from male fetuses (p=0.010; Figure 4D and E) and in the brains of both male and female fetuses, relative to the saline treated controls (p=0.042; p=0.040; Figure 5E). The combination of both oxy and melatonin significantly increased eIF2α phosphorylation (p=0.011; Figure 4E) and the p-eIF2α:eIF2α ratio in placentas from both male and female fetuses, compared to controls (p=0.032; p=0.041; Figure 4F). A similar response was observed in the fetal brains (p<0.05; Figure 5E, F). Collectively, these findings indicate that oxy induced activation of the ISR in both the placenta and fetal brain and crucially, melatonin co-treatment did not attenuate ISR activation in our rat model.

**Figure 4.**
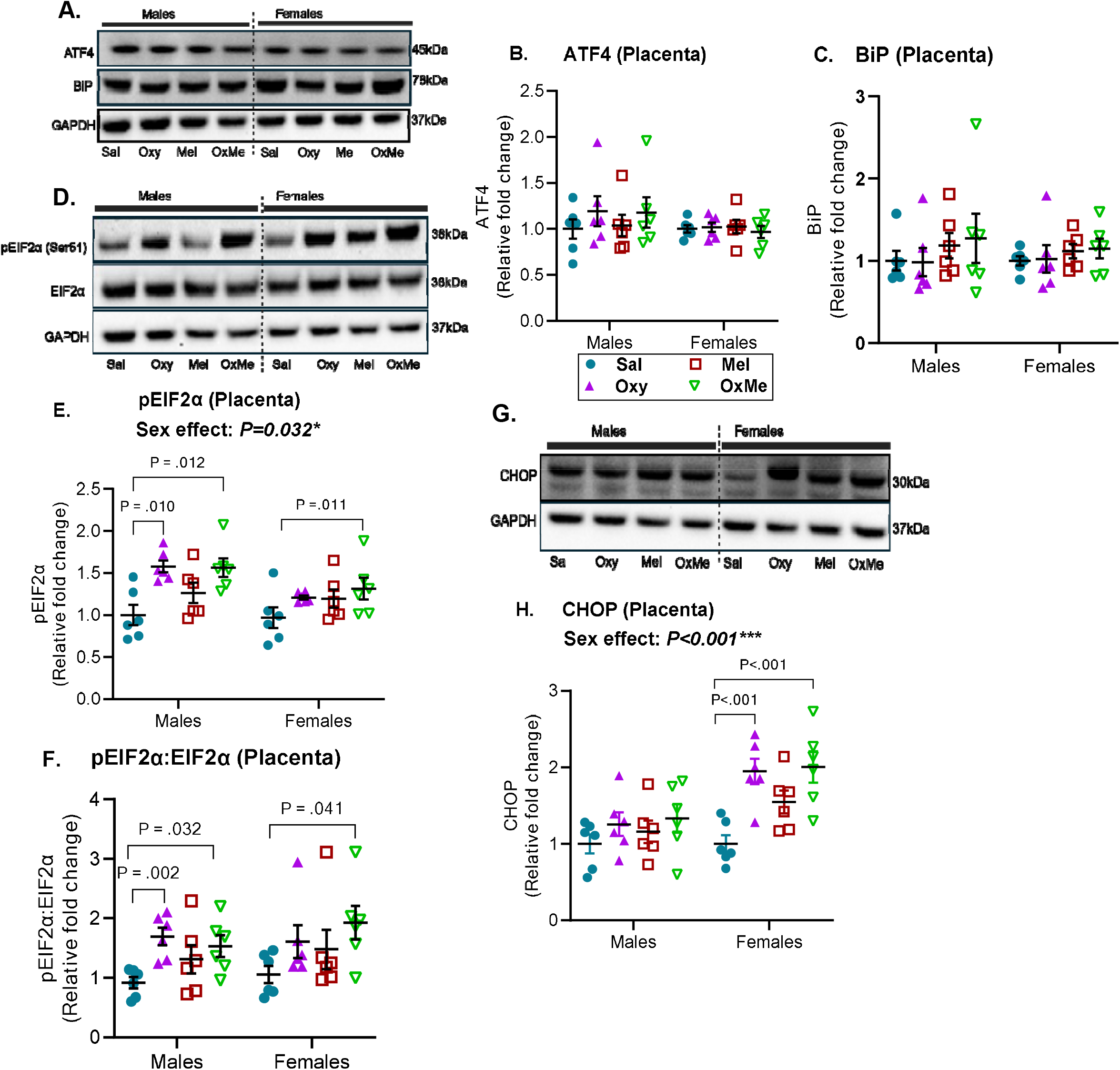
Effects of oxy and melatonin on integrated stress response markers in GD19.5 placenta. (A-C) Representative western blot images and densitometric quantification of ATF4 and BiP. (D-F) Representative western blot images, densitometric quantification of pEIF2a and graph of pEIF2a: EIF2a ratio. (G-H) Representative western blot images and densitometric quantification of CHOP. All samples were normalized to GAPDH. Data were analyzed by a 2-way ANOVA followed by Tukey’s post hoc test and presented as mean ± SEM. n=6/sex/group. *P*<0.05 was considered statistically significant.

**Figure 5.**
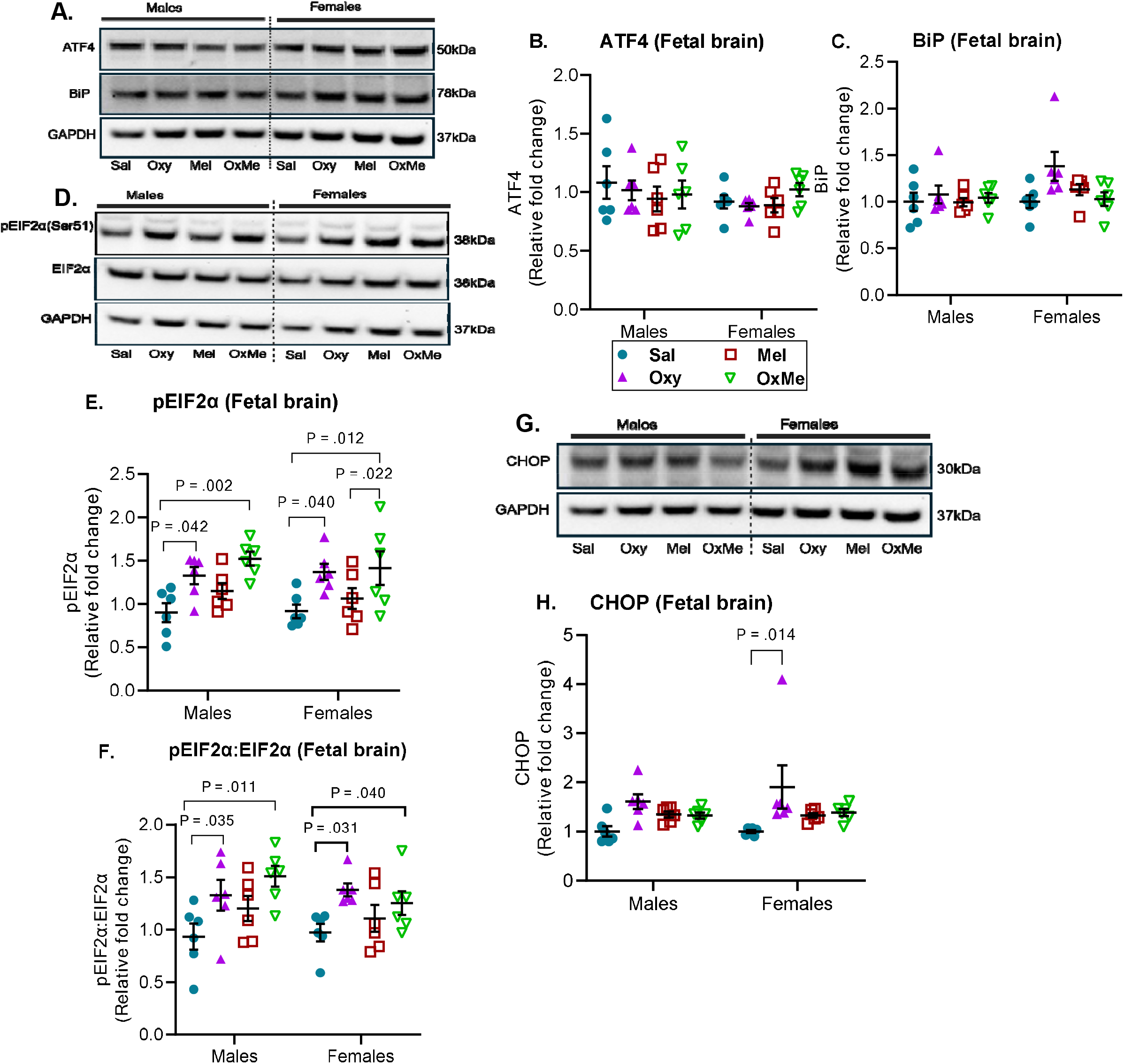
Effects of oxy and melatonin on integrated stress response markers in GD19.5 fetal brain. (A-C) Representative western blot images and densitometric quantification of ATF4 and BiP. (D-F) Representative western blot images and densitometric quantification of pEIF2a and EIF2a. (G-H) Representative western blot images and densitometric quantification of CHOP. All samples were normalized to GAPDH. Data were analyzed by a 2-way ANOVA followed by Tukey’s post hoc test and presented as mean ± SEM. n=6/sex/group. *P*<0.05 was considered statistically significant.

### Prenatal oxy exposure induced sex-specific CHOP expression in the placenta and fetal brain

We investigated expression levels of CHOP, as it can be upregulated through both ATF4-dependent and ATF4-independent mechanisms. CHOP expression was unaltered by oxy exposure or melatonin treatment in placentas and brains from male fetuses (Figure 4G,H; Figure 5G, H); however, CHOP expression was significantly higher in placentas from female fetuses exposed to oxy alone (p<0.001; Figure 4G,H), or oxy and melatonin (p<0.001), compared to controls. Additionally, oxy exposure modestly but significantly increased CHOP expression in the brains of female fetuses (p=0.014; Figure 5H). These findings identify tissue-specific and sex-dependent pattern of downstream ISR signaling in response to oxy exposure.

### Oxy promotes apoptotic signaling in the fetal brain, which is partially mitigated by melatonin treatment

To assess whether the observed activation of the ISR and CHOP translated to enhanced apoptotic signaling in the placenta and fetal brain, the expression levels of cleaved caspase-8, cleaved caspase-9, and cleaved caspase-3 were quantified. In the placenta, there was no difference in the expression level of cleaved caspase-3 across all the groups (Figure 6A, B). In the fetal brain, expression of cleaved caspase-8 was not altered by oxy exposure or melatonin treatment, indicating that the extrinsic apoptotic signaling pathway was not activated in our rat model (Figure 6C, D). However, levels of cleaved caspase-9 were significantly elevated in the brains of both male (p<0.001; Figure 6E) and female fetuses (p=0.023; Figure 6E) exposed to oxy. In males, melatonin co-treatment significantly reduced the levels of cleaved caspase-9 to control levels (p=0.020; Figure 6E). This protective effect was not observed in the brains of female fetuses (Figure 6E). However, despite the upstream attenuation of caspase-9 activation, cleaved caspase-3 expression was significantly elevated by oxy exposure in the brains of both male and female fetuses (p=0.003; p=0.014; Figure 6B). Melatonin co-treatment offered no protection against the pro-apoptotic effects of oxy. These findings indicate that oxy exposure activates the intrinsic apoptotic pathway in the fetal brain of both sexes, but this is not inhibited by melatonin.

**Figure 6.**
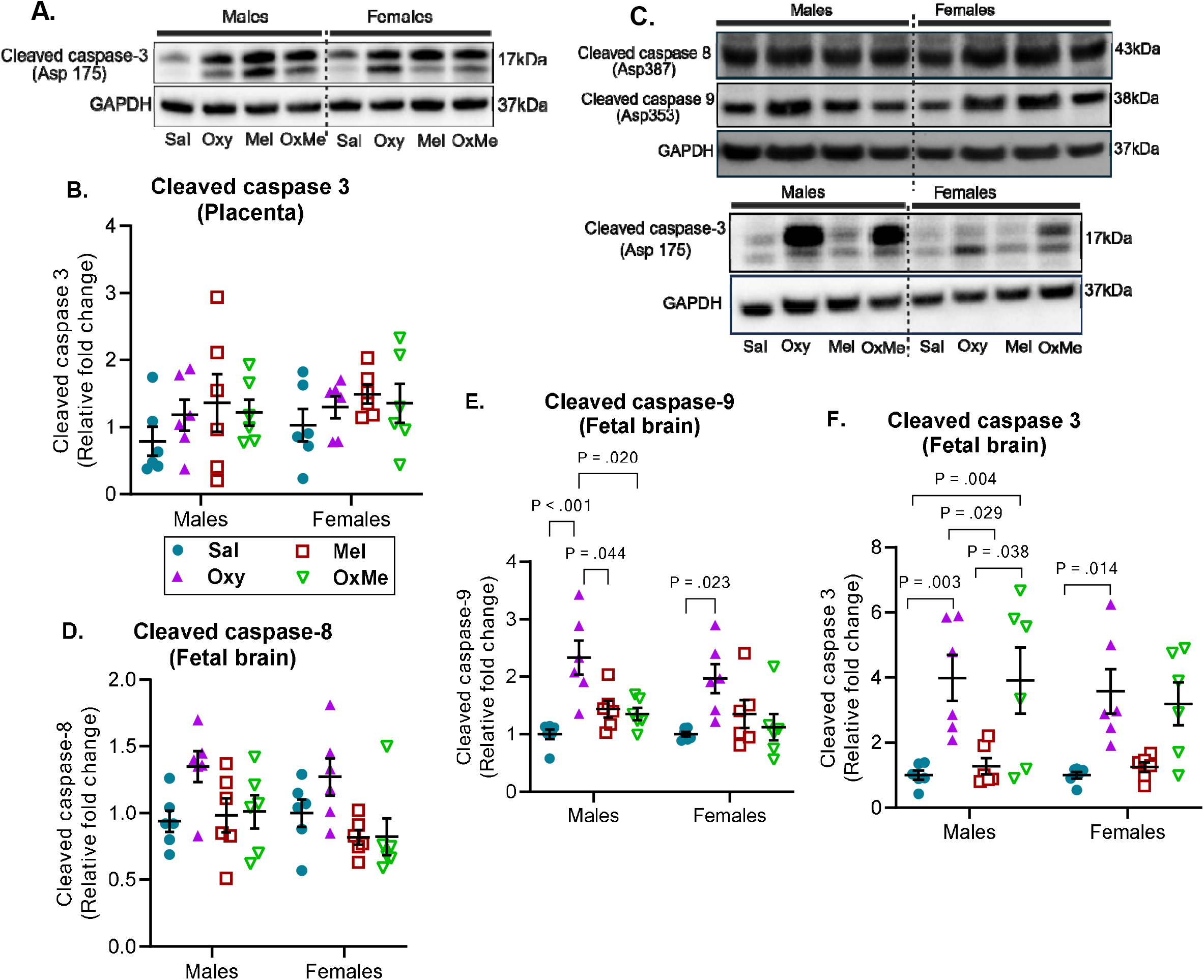
Effects of oxy and melatonin on integrated stress response markers in GD19.5 fetal brain. (A-C) Representative western blot images and densitometric quantification of ATF4 and BiP. (D-F) Representative western blot images and densitometric quantification of pEIF2a and EIF2a. (G-H) Representative western blot images and densitometric quantification of CHOP. All samples were normalized to GAPDH. Data were analyzed by a 2-way ANOVA followed by Tukey’s post hoc test and presented as mean ± SEM. n=6/sex/group. *P*<0.05 was considered statistically significant.

## DISCUSSION

Prenatal opioid exposure represents a significant public health challenge that causes adverse pregnancy and fetal outcomes, yet the molecular consequences in the placenta and fetal brain are not completely understood. Using a rat model of chronic, prenatal oxy exposure, we investigated whether oxidative stress, antioxidant defenses, placental inflammation, ER stress, and downstream apoptotic signaling would be induced by oxy in either the placenta or fetal brain and whether melatonin co-treatment would mitigate any adverse response.

Although opioid-induced oxidative stress has been described in the blood of addicted patients^47^, neuronal and microglial cell lines^48, 49^, we did not observe any increase in lipid peroxidation markers or antioxidant response in either the placenta or fetal brain. This is consistent with reports suggesting that opioid-induced oxidative stress is dependent on the specific chemical properties of the opioid ligand, the dosing threshold, tissue type, developmental stage differences, and metabolic state^50, 51^. Moreover, the differences in opioid-induced oxidative imbalances differ by opioid class, for instance, among patients on chronic opioid therapy, oxy was associated with greater oxidative stress in plasma compared to buprenorphine; however, oxy also elicited a higher antioxidant response, suggesting that there could be an adaptive response to oxy-induced oxidative burden^52^. Additionally, tissue-specific expression of antioxidants and metabolic enzymes may contribute to differential oxidative stress responses across tissues^28, 29, 50, 53^. These factors may explain why some studies report significant oxidative stress in response to opioid exposure, while others do not, potentially explaining the absence of oxidative damage or upregulation of antioxidant response genes in our model. Given that the KEAP-1/NRF2 axis is typically activated in response to an oxidative insult as a compensatory defense mechanism^54, 55^, the absence of NRF2 pathway activation in our present study further supports our findings that neither oxy nor melatonin induced oxidative stress at this gestational time point. While we cannot exclude that oxidative stress might have occurred at earlier gestational time points, these findings redirect our focus toward other stress pathways that can be activated independently of ROS accumulation.

Despite the absence of oxidative stress, oxy induced a selective inflammatory response, manifested as elevated IL-1β expression in both male and female placentas, while the expression of TLR4 and TNF-α were unchanged. IL-1β maturation and secretion are primarily driven by inflammasome activation, usually via NLRP3, which can be triggered by a range of sterile stimuli, including metabolic byproducts and cellular stress signals^56^. Given that levels of TNF-α, which is directly downstream of TLR4/NF-κB signaling^57, 58^, were unaffected in our study, this suggests that the source of IL-1β elevation is likely not TLR4-driven. Indeed, oxy-induced upregulation of IL-1β expression may be mediated via activation of the NLRP3 inflammasome, which has previously been linked to opioid receptor-mediated signaling^59, 60^. Additionally, we observed that melatonin treatment reduced mean placental IL-1β expression by 40%, which is consistent with findings from other studies that reported partial attenuating effects of melatonin on proinflammatory cytokines^61, 62^.

Our present study also highlights the dissociation between ISR activation and upregulation of canonical ER stress in response to chronic oxy exposure. Expression of BiP, the master ER chaperone and primary upstream regulator of the UPR, was unaffected in both placenta and fetal brain, indicating that the canonical UPR was not triggered^63, 64^. Despite this, eIF2α phosphorylation was significantly elevated in the placenta in a sex-specific manner, with males showing higher levels than females. Similarly, in fetal brain, eIF2α phosphorylation was significantly increased. eIF2α can be phosphorylated by at least four kinases such as PERK, HRI, PKR, and GCN2^32, 33, 65^. Given that PERK activation typically occurs concurrently with BiP induction as part of the broader UPR, the absence of BiP elevation in the context of elevated p-eIF2α in our rat model points toward a non-PERK kinase activation mechanism. This pattern is consistent with reports of increased eIF2α phosphorylation in the absence of BiP induction in the brain stem of oxy-treated rats^14^. Moreover, elevated levels of placental IL-1β induced by oxy exposure could also directly or indirectly activate HRI, PKR, and GCN2, driving eIF2α phosphorylation and ISR activation independently of PERK^66-68^.

Despite increased phosphorylation of eIF2α in both placenta and fetal brain, levels of ATF4, the transcription factor canonically induced downstream of p-eIF2α, were unchanged in our rat model. This is consistent with the findings of several studies which reported that elevated p-eIF2α does not inevitably lead to ATF4 protein accumulation, and that the relationship between p-eIF2α signaling and downstream ATF4 expression is highly context-dependent, varying with the nature of the upstream stress stimulus and the cellular environment^69, 70^.These findings could be attributed to the fact that ATF4 is short-lived and subject to rapid proteasomal degradation^71, 72^, which necessitates continuous and active translational input to maintain detectable steady-state levels, thus any transient accumulation could be missed at a single collection time point. Additionally, p-eIF2α-mediated ATF4 induction is not guaranteed, as the nature, duration, and context of the upstream stress stimulus can determine whether global translational repression overrides the mechanism by which ATF4 mRNA escapes that shutdown and gets selectively translated^69, 73, 74^ such that increased p-eIF2α can occur in the absence of detectable ATF4 accumulation depending on the stressor type and availability of ATF4 mRNA.

CHOP is a pro-apoptotic transcription factor frequently induced as a consequence of sustained ISR and is considered a marker of terminal stress signaling^65, 75^. In our model, we observed sex differences in CHOP expression in both the placenta and fetal brain. Prenatal oxy exposure significantly increased CHOP expression in the placentas and brains of female fetuses, while male tissues were unaffected. The elevation of CHOP in the context of unchanged ATF4 is mechanistically noteworthy, as CHOP can be induced through ATF4-independent mechanisms, including NF-κB, p38 MAPK, and other stress-activated signaling cascades^76, 77^. Whether the female-specific CHOP induction reflects differential upstream kinase activity, sex-dependent transcriptional regulation, or hormonal modulation of stress pathway is unknown from the findings our study. However, sex differences in placental stress responses have been well documented, though directionality appears to be context-and-stressor-dependent^78^ with female placentas generally mounting significant apoptotic responses under adverse gestational conditions in the context of prenatal opioid exposure^4, 79^; our findings are consistent with that pattern.

Given that CHOP is a downstream component of the ISR pathway, and its upregulation does not necessarily imply apoptotic commitment, it may also represent an adaptive response to cellular stress^33, 63, 70^, therefore we investigated the expression levels of cleaved caspase-8, 9 and 3. In the placenta, expression levels of cleaved caspase-3 were not altered regardless of fetal sex or treatment, suggesting that despite IL-1β elevation and ISR activation, the placenta is relatively resistant to pro-apoptotic stimuli at this gestational time point. This could be because of the intrinsic placental buffering mechanisms that could prevent downstream caspase-3 activation^80-82^. Trophoblast cells have been reported to constitutively express endogenous anti-apoptotic molecules including inhibitor of apoptosis (IAP) such as XIAP, cIAP-1, and cIAP-2, which act downstream of pro-apoptotic signals to suppress effector caspase activation. These molecules prevent further propagation of death signals even when upstream caspase cascades are activated^80-82^. In the fetal brain, levels of cleaved caspase-8 were not altered by oxy exposure, thus ruling out the activation of the extrinsic, cell death receptor-mediated apoptotic pathway^83^. In contrast, oxy exposure induced increased levels of cleaved caspase-9 in the fetal brains of both sexes, indicating the activation of the intrinsic mitochondria-dependent apoptotic pathway. In males, melatonin significantly reduced caspase-9 cleavage, which is consistent with the known mitochondrial protective properties of melatonin^36^ and suggests a plausible mechanism of action at the level of mitochondrial membrane integrity. However, this upstream attenuation did not translate into reduced levels of cleaved caspase-3 in the fetal brain, indicating that caspase-3 activation was maintained despite reduced levels of cleaved caspase-9. This could reflect caspase-3 amplification through alternative pathways or incomplete inhibition of the apoptosome. Collectively, we observed greater apoptotic vulnerability in the oxy-exposed fetal brain compared to the placenta, in line with previous studies describing the low apoptotic threshold of the developing brain to opioid exposure and intrauterine stress^84-86^. Additionally, we demonstrated that the ability of melatonin to attenuate the observed oxy-induced upregulation of caspase activation may be sex-dependent.

We acknowledge that our present study has some limitations that will require further investigations. Placental and brain tissues were collected at a single time point, thus limiting our ability to study variations in signaling responses that occur throughout gestation. Additional tissue harvests in early and mid-pregnancy will provide insight into any stage-specific responses to oxy-induced stress signaling. Also, melatonin was administered at a single dose and at a fixed zeitgeber time, so whether the observed effects reflect dose-dependent responses, circadian phase-specific sensitivity, or both remains unclear. Future studies should examine multiple doses across different zeitgeber times to tease apart these variables. Expression levels of kinases upstream of eIF2α, which mediate its phosphorylation were not measured, and the mechanistic basis for the IL-1β elevation cannot be definitively attributed to a specific stress signaling pathway without investigating involvement of the NLRP3 inflammasome or NF-κB signaling. Complementary IHC analysis of stress-related markers in brain tissue sections would further corroborate the molecular findings reported here and is planned for future work. Finally, our study is limited to prenatal observations, and the postnatal consequences of the placental and fetal brain changes reported here, including potential neurodevelopmental outcomes, is planned for future work.

## CONCLUSIONS

Despite these limitations, our findings demonstrate that prenatal oxy exposure induces selective placental IL-1β elevation, activates the ISR in both the placenta and fetal brain, and activates the intrinsic apoptotic pathway in the fetal brain, without triggering canonical ER stress or antioxidant defenses. The downstream consequences of ISR activation are in part sex-specific, and while melatonin co-treatment provided partial mitigation of some markers of stress and apoptotic signaling, it did not broadly attenuate ISR induction or cleaved caspase-activation. These findings advance our understanding of the tissue-specific and sex-dependent responses to prenatal oxy exposure and underscore the need for mechanistic follow-up studies to identify intervention targets.

## Author Contributions

**IOA** - Data curation, Formal analysis, Investigation, Project administration, Validation, Visualization, Writing – original draft, Writing – review & editing. **KK** – Investigation, Validation, Visualization, Writing – original draft, Writing – review & editing. **HMK** - Investigation, Project administration, Validation, Writing – review & editing. **PNM** - Investigation, Validation, Writing – review & editing. **COA** - Investigation, Validation, Writing – review & editing. **VLS** - Investigation, Project administration, Writing – review & editing. **JSD** - Conceptualization, Funding acquisition, Resources, Writing – review & editing. **ESP** - Conceptualization, Funding acquisition, Resources, Writing – review & editing. **GP** - Conceptualization, Funding acquisition, Methodology, Resources, Supervision, Writing – review & editing. **LKH** - Conceptualization, Funding acquisition, Investigation, Methodology, Project administration, Resources, Supervision, Writing – original draft, Writing – review & editing.

